# ATP hydrolysis-driven structural transitions within the *S. cerevisiae* Rad51 and Dmc1 nucleoprotein filaments

**DOI:** 10.1101/2025.03.19.644215

**Authors:** Yeonoh Shin, Stefan Y. Kim, Eric C. Greene

**Affiliations:** Department of Biochemistry & Molecular Biophysics, Columbia University Irving Medical Center, New York, NY, 10032, USA

## Abstract

Homologous recombination (HR) is essential for the maintenance of genome stability and for generating genetic diversity during meiosis. The eukaryotic protein Rad51 is member of the Rad51/RecA family of DNA recombinases and is responsible for guiding the DNA pairing reactions that take place in HR during mitosis. Dmc1 is a meiosis-specific paralog of Rad51 and is responsible for the DNA pairing reactions that take place in HR during meiosis. Rad51 and Dmc1 are both ATP-dependent DNA-binding proteins and both form extended helical filaments on ssDNA which are key intermediates in HR. The stability of these nucleoprotein filaments is highly regulated and is also tightly coupled to nucleotide binding and hydrolysis. ATP binding promotes filament assembly whereas the hydrolysis of ATP to ADP reduces filament stability to promote filament disassembly. Here, we present CryoEM structures of the *Saccharomyces cerevisiae* recombinases Rad51 and Dmc1 in the ADP-bound states and provide a detailed structural comparison to the ATP-bound filaments. Our findings yield insights into the structural transitions that take place during the hydrolysis of ATP to ADP and suggest a new model for how these structural changes may be linked to nucleoprotein filament disassembly.

## INTRODUCTION

Homologous recombination (HR) is essential for the maintenance of genome integrity and is a major driving force in genome evolution [1, 2]. HR plays important roles in double-stranded DNA break (DSB) repair [3, 4], the rescue of stalled or collapsed replication forks [5, 6], chromosomal rearrangements [7–9], horizontal gene transfer [10], and meiosis [11–13]. The protein participants, nucleoprotein structures, and general HR reaction mechanisms are broadly conserved among all kingdoms of life [14–17].

The DNA pairing reactions that take place in HR are promoted by the Rad51/RecA family of DNA recombinases, which are ATP–dependent proteins that form extended helical filaments on single stranded DNA, that are often referred to as presynaptic complexes [14–17]. During HR, a single–stranded DNA (ssDNA) serves as a platform for the assembly of the presynaptic complex which then pairs the bound ssDNA with the complementary strand from a homologous double–stranded DNA (dsDNA) from elsewhere in the genome, resulting in displacement of the non–complementary strand [3, 14–16]. These displacement loop (D–loop) intermediates can then be channeled through several mechanistically distinct pathways to restore the continuity of the originally damaged DNA [3, 14–16].

Most eukaryotes have two Rad51/RecA recombinases: Rad51, which is expressed constitutively, and Dmc1, which is only expressed in meiosis [11, 13, 18–20]. Rad51 and Dmc1 are thought to have arisen from an ancient gene duplication event, and the emergence of Dmc1 as a separate lineage may have coincided with the emergence of meiosis and sexual reproduction [21–24]. These two proteins remain closely related; for instance, *S. cerevisiae* Rad51 and Dmc1 share 45% sequence identity and 56% sequence similarity and both proteins perform the same basic biochemical function, namely the pairing of homologous DNA sequences [11, 13]. Rad51 and Dmc1 were identified over 25 years ago [18, 25], yet we still do not fully understand of why most eukaryotes require two recombinases [11, 13]. Prevailing hypotheses are that (*i*) each recombinase must interact with a specific subset of mitotic- or meiotic-specific accessory factors, (*ii*) there are biochemical differences between the recombinases making them uniquely suited to mitotic or meiotic HR [11, 13]. There are examples of Rad51- and Dmc1-specific interacting factors [11, 13], and Rad51 and Dmc1 respond differently when presented with mismatch–containing HR intermediates [26–28], suggesting that aspects of both hypotheses may be correct.

It has long been known that DNA within the active ATP-bound presynaptic complex is stretched by approximately 50% relative to B-form DNA and crystal structures of RecA–ssDNA presynaptic and RecA–dsDNA postsynaptic complexes reveal that the DNA is organized into near B–form base triplets separated by ∼8 Å between adjacent triplets [29]. This base triplet structural organization likely underpins homology recognition and the ability of the Rad51/RecA family of recombinases to promote DNA strand invasion in 3–nt steps [29–33]. This structural organization of the DNA now recognized to be a conserved feature of the nucleoprotein filaments made by Rad51/RecA family members.

Presynaptic complex assembly and disassembly are linked to the ATP hydrolysis cycle [14, 34]. Presynaptic complex assembly begins with an initial nucleation event with the recombinases in the high affinity ATP-bound state, followed by elongation as new ATP-bound protomers are added to the ends of the growing filaments [35–38]. Disassembly also takes place from the filament ends as the recombinase protomers hydrolyze ATP and are converted to the weakly bound ADP-state [39]. Based upon early studies with bacterial RecA, it is thought that the transition between the ATP- and ADP-bound states is also linked to changes in filament structure and function. Filaments in the ATP-bound state are more elongated with a helical pitch ranging between approximately 90 Å and 130 Å, while filaments in the ADP-bound state are thought to be more compressed with a helical pitch ranging from approximately 60 Å to 80 Å [29, 40–43]. Given the link between the nucleotide-bound state and the structure and function of the presynaptic complex, these transitions represent a potentially important regulatory control mechanism.

We have previously reported the CryoEM structures for the ATP-bound states of *S. cerevisiae* Rad51 and Dmc1, which correspond to the highly elongated states expected for biochemically active nucleoprotein filaments [44]. Here, we present CryoEM structures of *S. cerevisiae* Rad51 and Dmc1 nucleoprotein filaments in their ADP-bound states. Comparison of the ATP-bound and ADP-bound states show that the nucleoprotein filaments undergo a large structural transition resulting in weakened contacts between proteins within the nucleoprotein filaments and disordered electron density for the ssDNA. However, instead of undergoing compression, as expected based on studies of bacterial RecA [40], we find that the ADP-bound states of both Rad51 and Dmc1 were even more highly elongated, consistent with studies of human RAD51 [45, 46], suggesting that the observed structural transitions may be broadly conserved among the eukaryotic members of the Rad51/Dmc1 family members but are distinct from bacterial RecA. We suggest a model in which the extension of the Rad51 and Dmc1 nucleoprotein filaments upon ATP hydrolysis would require increased tension within the bound ssDNA to maintain proper register with the protein filament. This effect, together with weaker protein-protein contacts, may help to explain how the ATP hydrolysis cycle is linked to Rad51 and Dmc1 nucleoprotein filament disassembly.

## RESULTS

### CryoEM structure of Rad51 and Dmc1 filaments in the presence of ADP

To solve the CryoEM structures of *S. cerevisiae* Rad51 and Dmc1 in the ADP-bound states we mixed the purified proteins with 96-nucleotide ssDNA oligonucleotide in the presence of 2 mM ADP. Under these conditions, Rad51 yielded filaments with a resolution of 3.37 Å (Figure 1A-B, Figure S1, Figure S3A-SB & Table S1). Interestingly, in contrast to our observations with Rad51, the Dmc1 filaments were interdigitated (Figure S2), similar to what was recently observed for human RAD51 filaments in the presence of ADP [46]. However, the only potential point of contact between the filaments was between residues E62 and R301, but the density of R301 is poor, suggesting that this does not represent a true interaction. At this time, we do not have any reason to believe that these interdigitated structures reflect a biologically meaningful intermediate. Therefore, the interdigitated filaments were computationally deconvolved into single helical filaments (Figure S2) yielding well defined structures with an overall resolution of 2.7 Å (Figure 1C-D, Figure S2, Figure S3C-S3D & Table S1).

**Figure 1.**
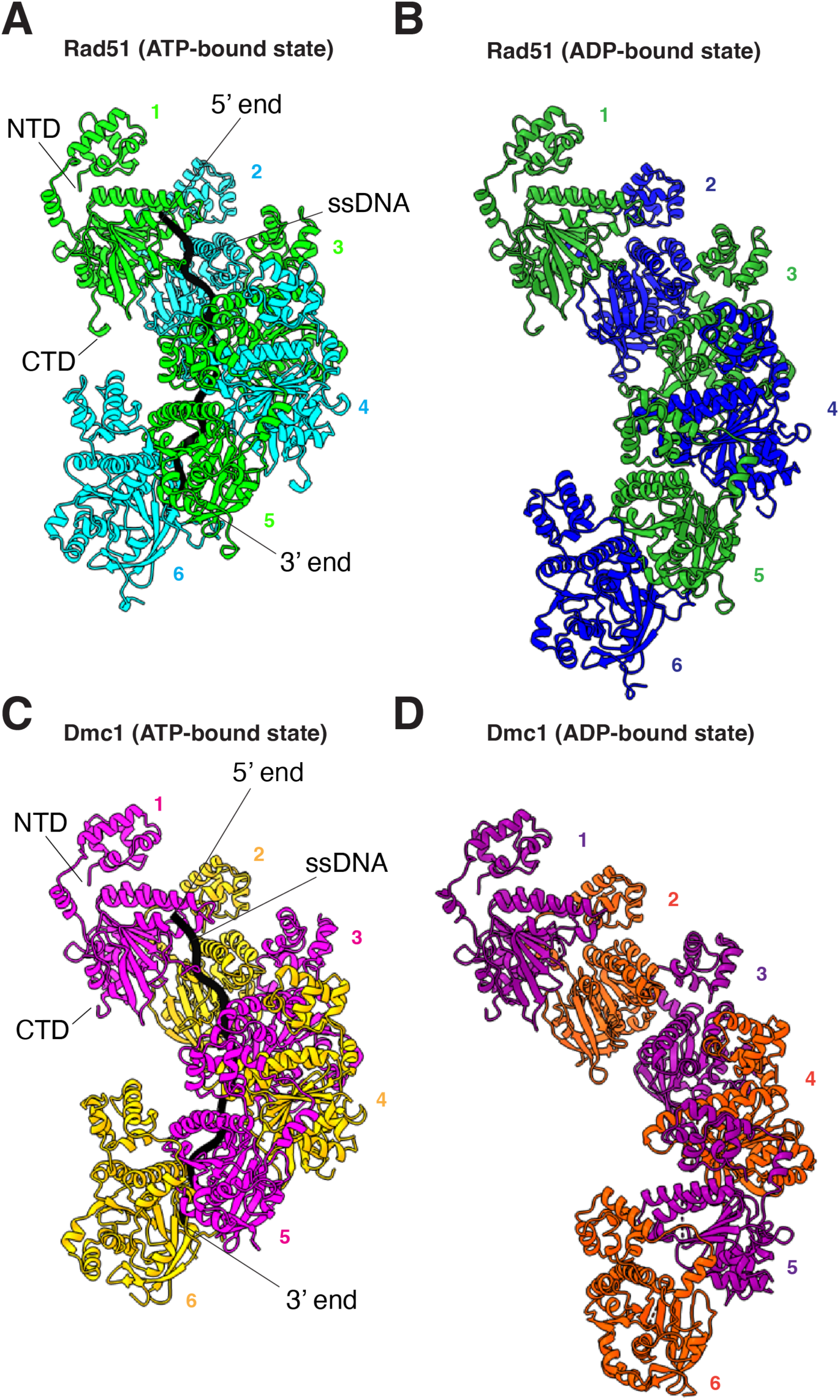
Overview of Rad51 and Dmc1 filament structures in the ADP-bound states. **(A)** Our previously published CryoEM structure of *S. cerevisiae* Rad51 nucleoprotein filament in the ATP-bound state (PDB ID: 9D46)[44]. A subsection of the nucleoprotein comprised of six Rad51 monomers is shown, the different protein monomers are highlighted in alternating cyan and light green, and the bound ssDNA is shown in black. **(B)** CryoEM structure of *S. cerevisiae* Rad51 bound in the ADP-bound state (PDB ID: 9NJK). A subsection of the nucleoprotein comprised of six Rad51 monomers is shown, and the different protein monomers are highlighted in alternating blue and green; the ssDNA is disordered within the structure. **(C)** Our previously published CryoEM structure of *S. cerevisiae* Dmc1 nucleoprotein filament in the ATP-bound state (PDB ID: 9D4N)[44]. A subsection of the nucleoprotein comprised of six Dmc1 monomers is shown, the different protein monomers are highlighted in alternating magenta and orange, and the bound ssDNA is shown in black. **(D)** CryoEM structure of *S. cerevisiae* Dmc1 bound in the ADP-bound state (PDB ID: 9NJR). A subsection of the nucleoprotein comprised of six Dmc1 monomers is shown, and the different protein monomers are highlighted in alternating purple and dark orange; the ssDNA is disordered within the structure.

We have previously reported CryoEM structures for *S. cerevisiae* Rad51 and Dmc1 in the ATP-bound states (PBD IDs: 9D46 & 9D4N)[44], and here we use these structures as references for comparison to the ADP-bound states. There were extensive changes in the overall structures of the nucleoprotein filaments assembled in the ADP-bound states compared to the ATP-bound states (Figure 1 & Figure S3). Most notably, there was a dramatic increase in the overall contour length of the protein filaments in the ADP-bound states. In the case of Rad51, the ATP-bound filaments exhibited a helical pitch of 99 Å with 6.4 protein monomers per helical turn and 19.2 nucleotides per turn, each protomer was rotated 56.4° relative to its nearest neighbors, and the filament had a maximum width of 87.6 Å (Figure 1A). In the Rad51 ADP-bound structure, the helical pitch increased to 132 Å, corresponding to an overall increase in the contour length of 33.3%, there were 6.5 monomers per turn and each protomer was rotated 51.6 degrees relative to its neighbors, and the filament had a maximum width of 95.2 Å (Figure 1B). In the case of Dmc1, the ATP-bound filaments exhibited a helical pitch of 98.4 Å with 6.35 protein monomers per helical turn and 19.05 nucleotides per turn, each promoter was rotated 55.5 degrees relative to its neighbors, and the filament had a width of 87.6 Å (Figure 1C). In the Dmc1 ADP-bound structure, the helical pitch increased to 133 Å, corresponding to an overall increase in the contour length of 35.2%, there were 6.6 monomers per turn and each promoter was rotated 47.4 degrees relative to its neighbors, and the filament had a maximum width of 97.5 Å (Figure 1D). The extended pitch of these filaments closely resembled the 130 Å pitch originally observed for the crystal structure of *S. cerevisiae* Rad51 [41] as well as the extended pitches observed for more recent CryoEM structures of human RAD51 in the ADP-bound state [45, 46].

The extended helical pitches observed for the ADP-bound states of Rad51 and Dmc1 coincided with the disordered ssDNA density, consistent with a weakening of the protein-DNA contacts, as has been previously reported for human RAD51 [45, 46], strongly suggesting that this structural change is a conserved property of the eukaryotic members of the Rad51/RecA family (Figure 1B & Figure 1D). It should be noted that formation of the Rad51 and Dmc1 filaments in the ADP-bound state required inclusion of the ssDNA substrate in the sample mixture, so although electron density for the ssDNA is not well resolved in the CryoEM data, it seems likely that the ssDNA is present within the protein filaments.

### Organization of the nucleotide-binding pocket

Rad51/RecA family members have bipartite nucleotide binding pockets comprised of highly conserved amino acid side chains from two adjacent protomers [29, 41]. The Walker A and Walker B nucleotide binding motifs are contributed by a single protomer and reside on the 3’ facing side of each protomer (Figure 2). Our CryoEM structures of Rad51 and Dmc1 reveal well-defined density for the ATP and ADP molecules within the nucleotide binding clefts in the ATP-bound and ADP-bound states, respectively (Figure 2A,2B & 2D,2E).

**Figure 2.**
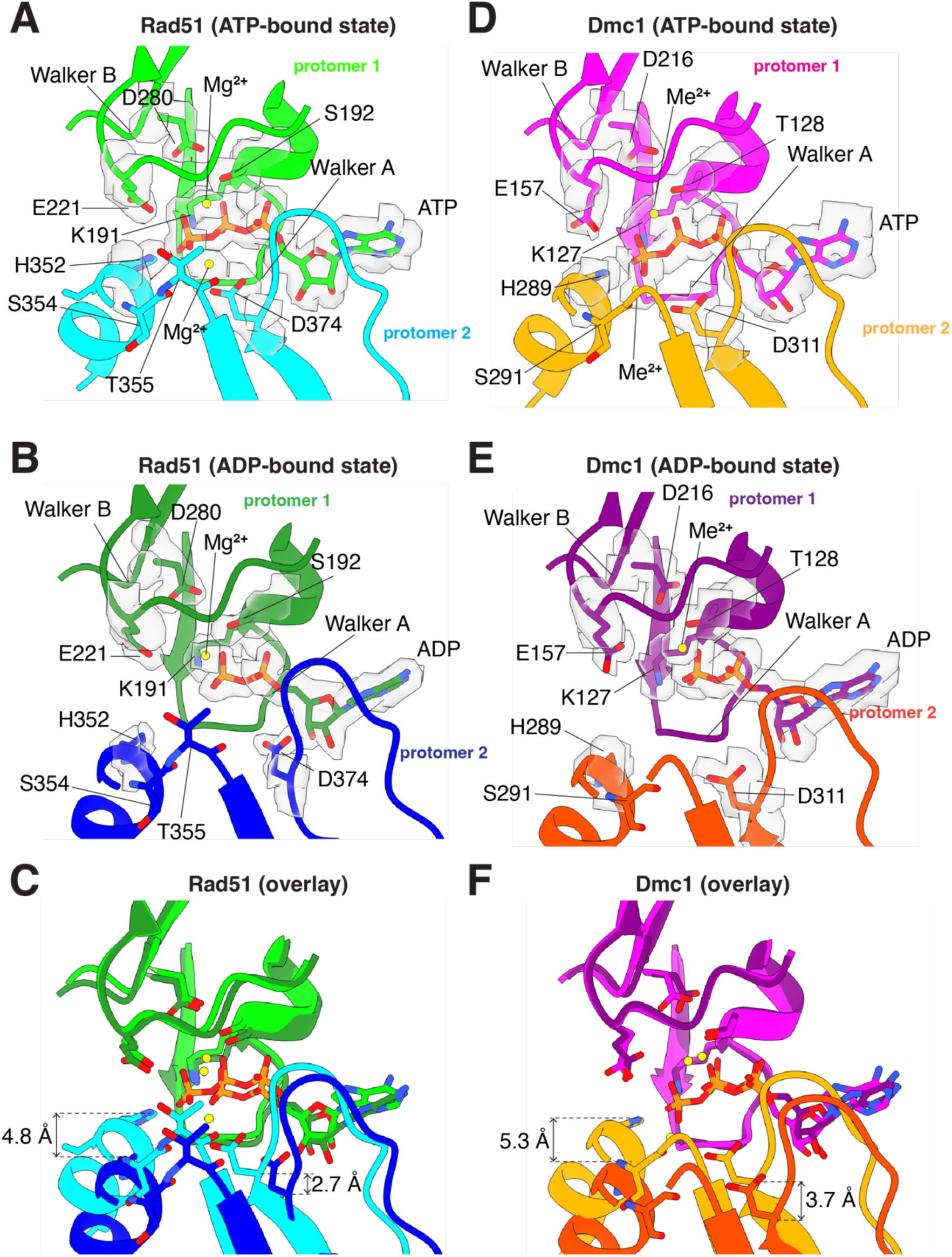
Comparison of the nucleotide-binding pockets in the ATP and ADP-bound states. **(A)** Nucleotide-binding pocket of *S. cerevisiae* Rad51 in the ATP-bound state. The Rad51 promoter in which the Walker A and Walker B motifs interact with the bound nucleotide is shown in light green and the second protomer is shown in cyan. The locations of important side chain residues and the two metal ions are indicated. **(B)** Nucleotide-binding pocket of *S. cerevisiae* Rad51 in the ADP-bound state. The Rad51 promoter in which the Walker A and Walker B motifs interact with the bound nucleotide is shown in green and the second protomer is shown in blue. The locations of important side chain residues and the two metal ions are indicated. **(C)** Overlay of Rad51 in the ATP- and ADP-bound states. **(D)** Nucleotide-binding pocket of *S. cerevisiae* Dmc1 in the ATP-bound state. The Dmc1 promoter in which the Walker A and Walker B motifs interact with the bound nucleotide is shown in magenta and the second protomer is shown in orange. The locations of important side chain residues and the two metal ions are indicated; here, the metal ions are indicated as Me2+ because we cannot distinguish between Mg^2+^ and Ca^2+^, both of which were present in samples with Dmc1 **(E)** Nucleotide-binding pocket of *S. cerevisiae* Dmc1 in the ADP-bound state. The Dmc1 promoter in which the Walker A and Walker B motifs interact with the bound nucleotide is shown in purple and the second protomer is shown in dark orange. The locations of important side chain residues and the two metal ions are indicated. **(F)** Overlay of Dmc1 in the ATP- and ADP-bound states.

There is a highly conserved histidine residue on the 5’ face of each recombinase protomer, H352 in Rad51 and H289 in Dmc1, both located with within α helix 14 (Figure S4), which interact in trans with the nucleotide binding pocket of the neighboring protomer and helps to coordinate the gamma phosphate of the bound ATP molecule (Figure 2A & Figure 2D). It has previously been suggested that these histidine residues act as sensors of ATP binding and help to coordinate communicate between adjacent recombinase protomers within the nucleoprotein filaments [47]. Previous studies have shown, mutation of H352 within *S. cerevisiae* Rad51 to either alanine, tyrosine, phenylalanine, lysine or methionine, severely compromises both ATP hydrolysis activity and nucleoprotein filament assembly at physiological pH [47]. Consistent with this model, we find that Rad51 H352 and Dmc1 H289 move away from the neighboring promoters when the nucleoprotein filaments are in the ADP-bound states, corresponding to a shift of 4.8 Å in the case of Rad51 and 5.3 Å for Dmc1 measured from the epsilon carbon if the histidine side chains (Figure 2C & 2F). Together with the previous study [47], this work further highlights the importance of H352 from Rad51 and the corresponding H289 from Dmc1 as important residues for sensing the nucleotide-bound state of the nucleoprotein filaments and because these interactions occur in trans they may contribute to allosteric communication within the nucleoprotein filaments.

For both Rad51 and Dmc1, we observe density for two metal ions within the nucleotide binding cleft in the ATP-bound states (Figure 2A & D). In the ADP-bound states, however, there is a loss of the second metal ion (Figure 2B & E), a phenomenon that has also been reported for human RAD51 [45]. For Rad51, the first metal ion is coordinated in cis by the sidechains of residues S192 and D280 which are located between the beta and gamma phosphate of the bound ATP molecule as well as residue E221 which is near the gamma phosphate group of the ATP (Figure 2A). In the case of Dmc1, the first metal ion is coordinated in cis by the sidechains of residues T128, D216 and E157 which are all positioned similarly to the equivalent residues from Rad51 (Figure 2D). These contacts with the first metal ion do not undergo any large changes in the ATP- versus ADP-bound states, for either Rad51 (Figure 2A-2C) or Dmc1 (Figure 2D-2F). The second metal ion is coordinated in trans by a highly conserved aspartic acid residue, D374 from Rad51 and D311 from Dmc1, from the 5’ facing side of the adjacent protomer (Figure 2A & 2D). The second metal ion also interacts in with the amide group from the peptide backbone of residues S354 and T355 from Rad51 and residue S291 from Dmc1 (Figure 2A & 2D). Comparison of the ATP-bound and ADP-bound states reveals that *S. cerevisiae* Rad51 D374 and Dmc1 D311 move away from the neighboring promoters in the ADP-bound states, corresponding to a distance (measured from the alpha carbon) of 2.7 Å in the case of Rad51 and 3.7 Å for Dmc1 (Figure 2A-2F). Importantly, this trans interaction with the second metal ion has also been reported for human RAD51 (D316) and archaeal RadA (D302), providing further evidence for the conservation of this mechanism [45, 48, 49].

Dmc1 requires the presence of Ca^2+^ for optimal *in vitro* activity [13, 50–52]. Although we see clear density for two metal ions in the nucleotide binding pocket of *S. cerevisiae* Rad51 and Dmc1, we cannot directly distinguish between Mg^2+^ and Ca^2+^ based on the electron density alone. In the case of Rad51, Mg^2+^ was the only divalent metal ion cofactor included in the filament assembly reactions, so it is reasonable to conclude that there are two Mg^2+^ ions bound within the Rad51 active site. However, for Dmc1, the filament assembly reactions required both Mg^2+^ and Ca^2+^, as has been well established in the literature [13, 50–52], so the bound metal ions could be Mg^2+^, Ca^2+^, or a mixture of both metal ions. Interestingly, superimposition of the Rad51 and Dmc1 structures from nucleoprotein filaments in the ATP-bound states reveals that the second divalent metal ion appears to be shifted by 2.1 Å in Rad51 compared to Dmc1 (Figure S5A-5C). Comparison of our *S. cerevisiae* Rad51 (determined with Mg^2+^ only) and Dmc1 (determined with Mg^2+^ and Ca^2+^) structures to a recent structure for human RAD51 in the ATP-bound state on ssDNA in the presence of only Ca^2+^ revealed that the second metal ion in the human RAD51 (PBD ID: 8BQ2; Figure S5D)[45] is shifted by 2.2 Å compared to the second metal ion in yeast Rad51 (Figure S5E), but matches well with the second metal ion found in yeast Dmc1 (Figure S5F). One possibility is that the differential positioning of the second metal ion in human RAD51 and *S. cerevisiae* Dmc1 reflects differential positioning due to the identity of the bound divalent metal ion, which might imply that Ca^2+^ is the second metal observed in our structure of *S. cerevisiae* Dmc1 even though Mg^2+^ was present in 13-fold molar excess over Ca^2+^ (20 mM MgCl_2_ versus 1.5 mM CaCl_2_).

### The FxxA polymerization motif remains intact in the ADP-bound state

Eukaryotic and archaeal members of the Rad51/RecA family of proteins harbor a conserved FxxA polymerization motif that is essential for nucleoprotein filament formation [25, 53]. This motif is found within the interdomain linker between the N-terminal five–helix bundle and core domain (Figure 3A-3B, and Figure S4) and is comprised of residues 144–FVTA–147 and 80–FIPA–83 for *S. cerevisiae* Rad51 and Dmc1, respectively (Figure 3C-3H). For Rad51, phenylalanine residue F144 binds within a hydrophobic pocket of the adjacent protomer formed by residues L216, I218, A248, A250, L261, A264, and M268 from the adjacent protomer, and A147fits into a hydrophobic pocket formed by F224, L244, and V247. Residue F224 is further stabilized by stacking interactions with Y249 and P226 in cis, and F150 in trans (Figure 3C). For Dmc1, phenylalanine residue F80 binds within a hydrophobic pocket of the adjacent protomer formed by residues A152, I154, S184, A186, L197, and L201 from the adjacent protomer, and residue A83 fits into the hydrophobic pocket formed by F160, L180, and V183. Residue F160 is further stabilized by stacking interactions with Y185 and P162 in cis and Q86 in trans (Figure 3F). Interestingly, the FxxA polymerization motifs remain fully intact upon transition to the ADP-bound state for both Rad51 and Dmc1, indicating that the disruption of this protein-protein interfacial contact does not directly contribute to nucleoprotein filament destabilization upon the hydrolysis of ATP to ADP (Figure 3E & 3H).

**Figure 3.**
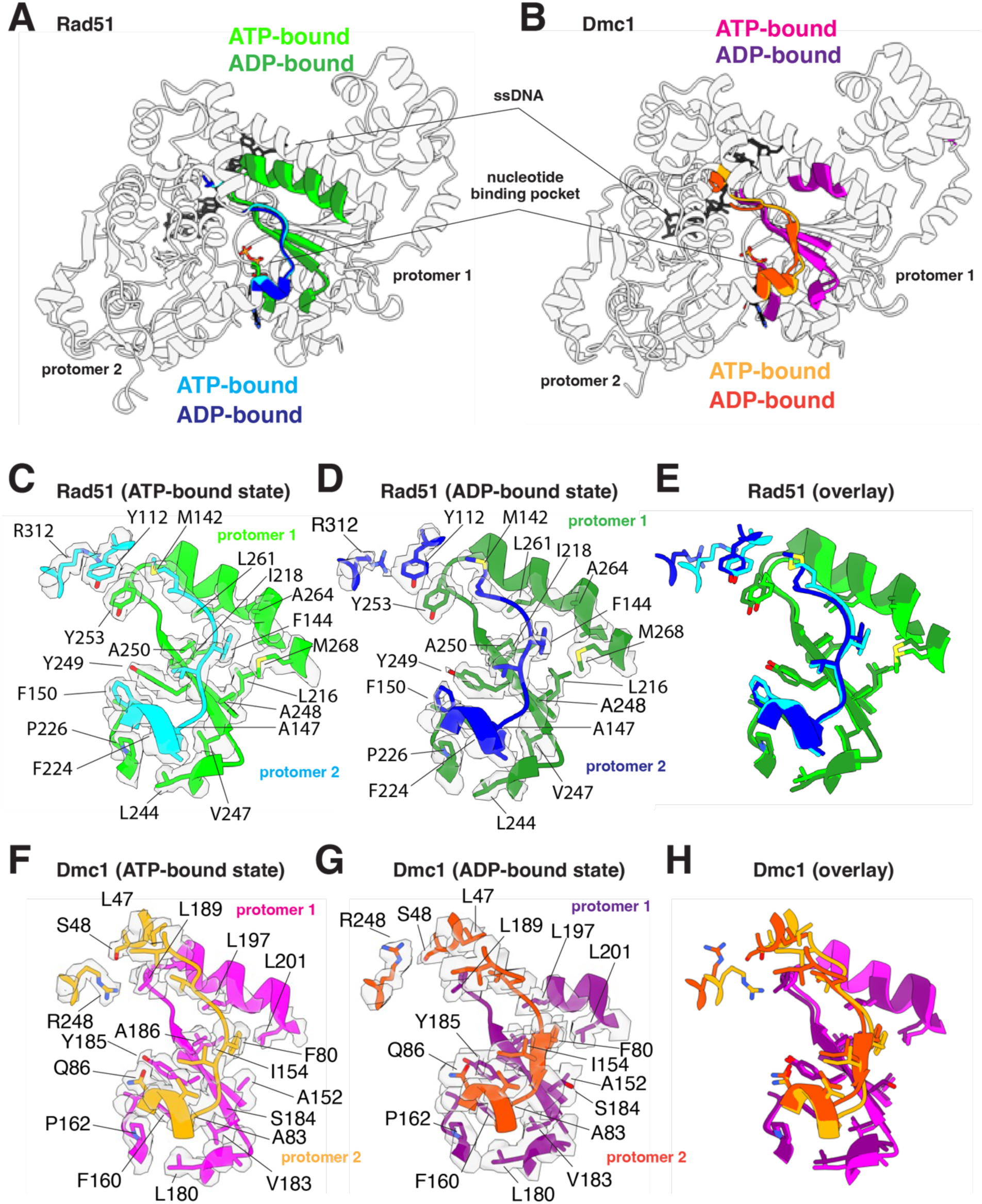
The FxxA polymerization interface remains largely unaltered in the ATP and ADP-bound states. **(A)** Overlay of two adjacent Rad51 protomers in the ATP- and ADP-bound states, as indicated, highlighting the location of the FxxA polymerization motif in one protomer and its binding cleft on the adjacent protomer. **(B)** Overlay of two adjacent Dmc1 protomers in the ATP- and ADP-bound states, as indicated, highlighting the location of the FxxA polymerization motif in one protomer and its binding cleft on the adjacent protomer. **(C)** Close-up view of the Rad51 FxxA polymerization motif interaction in the ATP-bound state. **(D)** Close-up view of the Rad51 FxxA polymerization motif interaction in the ADP-bound state. **(E)** Overlay of the Rad51 FxxA polymerization motif interaction in the ATP- and ADP-bound states. **(F)** Close-up view of the Dmc1 FxxA polymerization motif interaction in the ATP-bound state. **(G)** Close-up view of the Dmc1 FxxA polymerization motif interaction in the ADP-bound state. **(H)** Overlay of the Rad51 Dmc1 polymerization motif interaction in the ATP- and ADP-bound states.

### Disruption of a Dmc1-specific inter-protomer contact

In addition to the FxxA motif, Rad51 also harbors and highly conserved pair of stacked aromatic residues that is unique to the Rad51 lineage of the Rad51/RecA but absent in the Dmc1 lineage [44]. In *S. cerevisiae* Rad51, these correspond to amino acids residues Y112 and Y253, whereas the equivalent residues in *S. cerevisiae* Dmc1 are S48 and L189, which do not appear to interact with one another (Figure 3C & 3F). We have previously reported deep mutagenesis experiments which demonstrated that nonaromatic residues are largely not tolerated at these positions with *S. cerevisiae* Rad51, whereas many different residue pairs retain biological function in *S. cerevisiae* Dmc1 [44]. In contrast to the stacked aromatic residues at the Rad51 protein-protein interface, Dmc1 residues S48 and L189 do not appear to interact with one another, instead L189 appears to interact with residue R248 via van der Waals contacts which is stabilized by S48 through an intramolecular interaction, and residue L47 via hydrophobic interaction when in the ATP-bound state and this interaction is unique to Dmc1 (Figure 3F). The equivalent arginine residue in *S. cerevisiae* Rad51 is R312 which does not contact Y112 and Y253. The stacked Y112 and Y253 aromatic residues found in Rad51 are not disrupted upon transition to the ADP-bound state. In contrast, Dmc1 residue R248 shifts in the ADP-bound state and no longer interacts with residue L189 from the adjacent protomer. Thus, the disruption of this particular inter-protomer contact appears to be specific for Dmc1.

### Disruption of DNA-binding loop L1 and α helix 13 inter-protomer contacts

Comparison of the ATP- and ADP-bound structures for both Rad51 and Dmc1 reveal that transition to the ADP-bound state coincides with the rotation of a long α helix (helix 13; Figure S4) that is next to DNA-binding loop L1 and is involved in trans interactions with the L1 DNA-binding domain of the adjacent protomer (Figure 4A & 4B). Rotation of this helix coincides with the loss of DNA-binding loop L1 ssDNA contacts (Figure 4D & 4G and Figure S6) and also disrupts protein-protein contacts between adjacent protomers (Figure 4C-4H). In the case of Rad51, residues R308, R312 and D315 interact in *trans* with residues T288, D289, and R251 of the adjacent protomer in the ATP-bound state (Figure 4C). In addition, residue K216 from one promoter contacts residue E295 in the L1 DNA-binding loop of the adjacent protomer (Figure 4C). All of these contacts are disrupted in the ADP-bound state (Figure 4D & 4E). These interprotomer contacts are somewhat different in the case of Dmc1. For Dmc1, residues F244, R248, and E251 interact with V224 and R187 from the adjacent protomer in the ATP-bound state (Figure 4F). Lastly, similar to Rad51, Dmc1 residue R52 from one promoter contacts residue E231 in the L1 DNA-binding loop of the adjacent protomer (Figure 4C). As with Rad51, all of these contacts are disrupted in the ADP-bound state of Dmc1 (Figure 4G & 4H).

**Figure 4.**
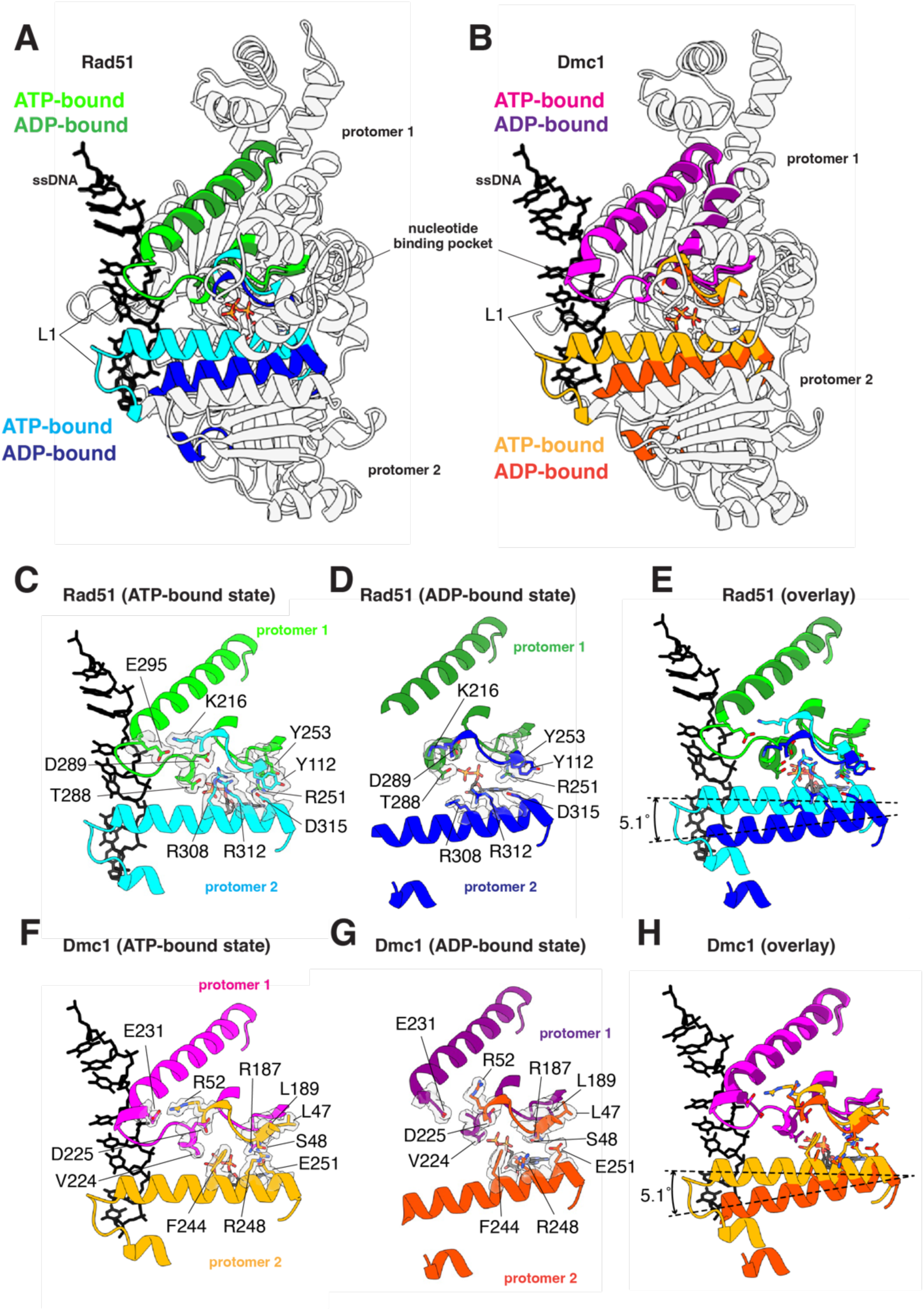
Loss of trans L1-L1 contacts between adjacent protomers in the ADP-bound state. **(A)** Overlay of two adjacent Rad51 protomers in the ATP- and ADP-bound states, as indicated, highlighting the location of the L1 DNA-binding loop connect to helix 13. **(B)** Overlay of two adjacent Dmc1 protomers in the ATP- and ADP-bound states, as indicated, highlighting the location of the L1 DNA-binding loop connect to helix 13. **(C)** Close-up view of the Rad51 *trans* interactions between L1 and helix 13 interaction in the ATP-bound state. **(D)** Close-up view of the Rad51 *trans* interactions between L1 and helix 13 interaction in the ADP-bound state. **(E)** Overlay of the Rad51 DNA-binding loop L1 and helix 13 in the ATP- and ADP-bound states. **(F)** Close-up view of the Dmc1 *trans* interactions between L1 and helix 13 interaction in the ATP-bound state. **(G)** Close-up view of the Dmc1 *trans* interactions between L1 and helix 13 interaction in the ADP-bound state. **(H)** Overlay of the Dmc1 DNA-binding loop L1 and helix 13 in the ATP- and ADP-bound states.

### Disruption of DNA-binding loop L2 structure and contacts

For both Rad51 and Dmc1 in the ATP-bound state, the L2 DNA-binding loop makes crucial contacts with the ssDNA and also makes protein-protein contacts between adjacent protomers within the nucleoprotein filaments (Figure 5). For Rad51 in the ATP-bound state, residues I345 and V328 form intramolecular hydrophobic interactions that helps to organize the overall shape of the L2 DNA-binding loop (Figure 5C). The peptide backbone of residue I345 also makes stabilizing intramolecular contacts with residue Q326 (Figure 5C) and residue P344 provides a structural kink point near the end of L2 allowing it to transit back towards the core of the protein (Figure 5C). Residues K343 and N348 both make contacts the ssDNA backbone and residue N348 also interacts in *trans* with residue K342 from the adjacent protomer (Figure 5C). Lastly, residue H352 interacts in *trans* with Q326, which is itself stabilized in *cis* with the side chains of E221 and R287, and the backbone of residue I345 (Figure 5C). These interactions are all weakened in the ADP-bound state as evidenced by the disordered electron density for both the ssDNA and many of the L2 amino acid residues (Figure 5D & 5E).

**Figure 5.**
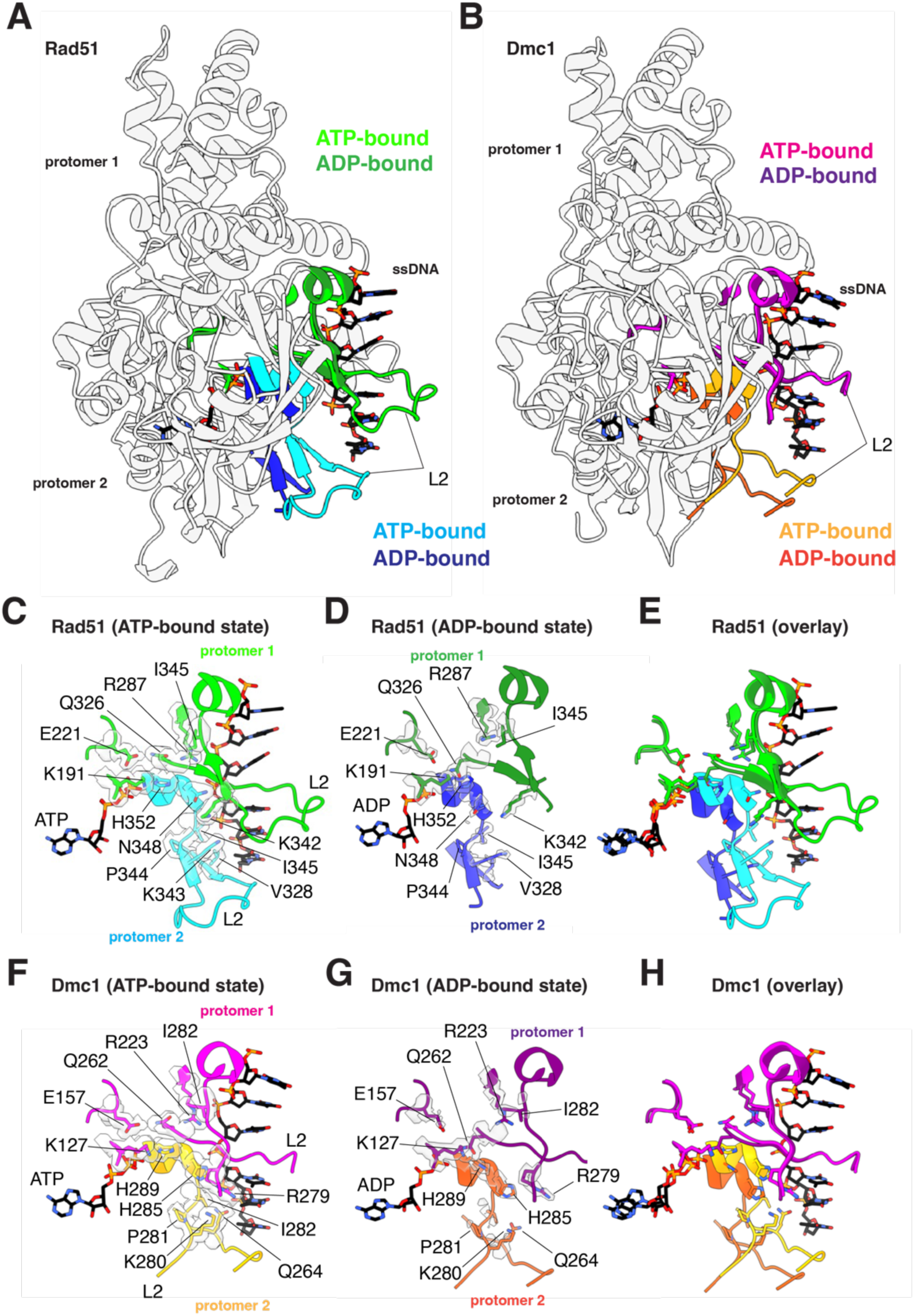
Loss of L2 ssDNA- and inter-protomer contacts in the ADP-bound state. **(A)** Overlay of two adjacent Rad51 protomers in the ATP- and ADP-bound states, as indicated, highlighting the location of the L2 DNA-binding loop, α helix 14, and β strands 6 and 7. **(B)** Overlay of two adjacent Dmc1 protomers in the ATP- and ADP-bound states, as indicated, highlighting the location of the L2 DNA-binding loop, α helix 14, and β strands 6 and 7. **(C)** Close-up view of the Rad51 L2 contacts with the bound ssDNA substrate and inter-protomer interactions in the ATP-bound state. **(D)** Close-up view showing the loss of Rad51 L2 contacts with the bound ssDNA substrate and inter-protomer interactions in the ADP-bound state. **(E)** Overlay of the Rad51 region encompassing the L2 DNA-binding loop, α helix 14, and β strands 6 and 7 in the ATP- and ADP-bound states. **(F)** Close-up view of the Dmc1 L2 contacts with the bound ssDNA substrate and inter-protomer interactions in the ATP-bound state. **(G)** Close-up view showing the loss of Dmc1 L2 contacts with the bound ssDNA substrate and inter-protomer interactions in the ADP-bound state. **(H)** Overlay of the Dmc1 region encompassing the L2 DNA-binding loop, α helix 14, and β strands 6 and 7 in the ATP- and ADP-bound states.

In the case of Dmc1, in the ATP-bound state, residues I282 and Q264 form intramolecular contacts between their side chains as well as contacts between the peptide backbone atoms which help to organize the overall structure of the L2 DNA binding loop (Figure 5F). The proline side chain of P281 provides a structural kink that allows L2 to transit back towards the core of the protein (Figure 5F). Residues H285, Q264, and K280 all make contacts with the ssDNA phosphate backbone (Figure 5F). Residues H289 and Q262 interact in *trans* between adjacent protomers, and the position of Q262 is organized by *cis* interactions with the side chains of residues E157 and R223, and the backbone of residue I282. Similar to Rad51, these interactions are all weakened in the ADP-bound state of Dmc1 as evidenced by the disordered density for both the ssDNA and many of the L2 amino acid residues (Figure 5G & 5H).

## DISUCUSSION

Our work shows that *S. cerevisiae* Rad51 and Dmc1 adopt a highly extended configuration when in the ADP-bound state. This large structural transition coincides with a weaking of DNA contacts, as evidenced as disordered electron density corresponding to the ssDNA and a weaking of contacts between adjacent protein subunits, supporting the hypothesis that ADP-bound state is an on path intermediate towards nucleoprotein filament disruption. These findings give insights into how the ATP hydrolysis cycle might be coupled to nucleoprotein filament disassembly and reveal interesting structural differences between the eukaryotic Rad51 and Dmc1 proteins compared to their bacterial homolog RecA, which may suggest that these proteins have distinct mechanistic approaches to regulating dissociation from ssDNA.

### Comparison between *S. cerevisiae* Rad51 and Dmc1

*S. cerevisiae* Rad51 and Dmc1 both form similar nucleoprotein filament structures in the presence of ADP and in each case the filaments are highly elongated compared to those formed in the ATP-bound state. These findings are in good agreement with previous studies which have shown that nucleoprotein filaments prepared with human RAD51 in the presence of ADP are also in a highly elongated state compared to those formed with ATP [45, 46]. Therefore, it is likely that this elongated ADP-bound state reflects a broadly conserved structural state for the eukaryotic Rad51 and Dmc1 recombinases.

It is notable that our structures of *S. cerevisiae* Rad51 and Dmc1 in the ADP-bound states bear a strong resemblance to the *S. cerevisiae* Rad51 crystal structure of *S. cerevisiae* Rad51, which represented a breakthrough in the field as it was the first reported high-resolution structure for Rad51 in a filament state [41]. Interestingly, the ssDNA and the DNA-binding loops were not visible within the Rad51 crystal structure, the filaments were prepared with ATPγS but no nucleotide cofactor was visible within the active site, perhaps because the ATPγS had undergone hydrolysis during crystallization, and instead a sulfate ion was visible within the active site [41]. Lastly, the helical pitch of 130 Å was observed in the *S. cerevisiae* Rad51 crystal structure [41], which closely corresponds to the helical pitch values of 132 Å and 133 Å that we report here for the ADP-bound states of *S. cerevisiae* Rad51 and Dmc1, respectively.

### Comparison of Rad51 and Dmc1 to bacterial RecA

Rad51/RecA family members are ATP-dependent DNA-binding proteins and the transition from the ATP- and ADP-bound states is linked to nucleoprotein filament disassembly [14, 34]. Thus, the ADP-bound forms of these nucleoprotein filaments should represent intermediates on the pathway towards protein dissociation. Based on structural studies of bacterial RecA, it had been thought that the active ATP-bound state was more elongated with a helical pitch ranging between approximately 90 Å and 130 Å, while inactive filaments in the ADP-bound state were thought to be more compressed with a helical pitch ranging from approximately 60 Å to 80 Å [29, 40–43]. Interestingly, based on the known high-resolution structures of eukaryotic recombinases in the ADP-bound states, including those presented here for *S. cerevisiae* Rad51 and Dmc1, and previous structures of human RAD51 [45, 46], the eukaryotic proteins may be behaving differently from bacterial RecA. Instead of transitioning to a compressed state, it appears as though Rad51 and Dmc1 undergo a structural conversion to a more elongated state which coincides with a loss of DNA contacts and disruption of contacts between the protomer-protomer interfaces. Indeed, we see no obvious evidence for the existence of compressed states for the ADP-bound nucleoprotein filaments for either Rad51 or Dmc1 under the conditions used for our sample preparation (Figure S1 & S2). These observations raise the possibly that Rad51 and Dmc1 might follow a different pathway towards DNA dissociation compared to the bacterial RecA protein. In this regard, it is interesting to note that although the core ATP-binding domains of eukaryotic Rad51 and Dmc1 and prokaryotic RecA are all strongly conserved, the protein-protein interfaces between protomers within the filaments are markedly different [40]. RecA has a C-terminal domain that contributes to the protein-protein interfaces and allosteric communication between adjacent proteins within a nucleoprotein filament, but this C-terminal domain is absent from the eukaryotic recombinases. Whereas Rad51 and Dmc1 have an N-terminal domain that is absent from RecA. In the case of Rad51 and Dmc1, the linker region between the N- and C-terminal domains contains a FxxA polymerization motif that contributes to inter-protomer contacts; however, this motif and the corresponding binding pocket appears to be absent from bacterial RecA. These differences in domain structure may be responsible for conferring distinct properties to the nucleoprotein filaments, which in this case manifest as structurally distinctive intermediates on the pathway towards nucleoprotein filament dissociation.

### Model for Rad51 & Dmc1 nucleoprotein filament disassembly

Rad51 and Dmc1 filaments in the high affinity ATP-bound state and bound to ssDNA that is held in a highly extended configuration. However, the extension is not isotropic, instead the base triplets are maintained in a near B-form DNA configuration, and the phosphodiester backbone between base triplets is highly extended which gives rise to the overall extension of the bound ssDNA, as originally reported for bacterial RecA [29]. In the case of *S. cerevisiae* Rad51, this corresponds to an axial rise of 7.2 – 7.3 Å between the last base of one triplet and the first base of the next triplet, and for *S. cerevisiae* Dmc1 these values are 7.0 – 7.2 Å (Figure 6A). Our work shows that the Rad51 and Dmc1 nucleoprotein filaments are even longer in the ADP bound state (Figure 6B). The ssDNA molecules within the ATP-bound states of the Rad51 and Dmc1 nucleoprotein filaments are approximately 1.6 times the length of an equivalent B-form dsDNA molecule. With respect to the ADP-bound states, if one were to consider a hypothetical scenario where the base triplets had to remain in register with the Rad51 or Dmc1 monomers in the same way as is observed in the ATP-bound structures, then the phosphodiester bonds between the base triplets would also have to increase up to ∼9.0 Å, which approaches the maximum extent to which the phosphodiester bond can be stretched and the bound ssDNA would have to be stretched to approximately 1.9 times the length of an equivalent B-form dsDNA (Figure 6B). If one assumes that the isotropic stretching forces experienced by a single stranded DNA molecule under tension by optical tweezers can be used to roughly approximate the forces experienced by the extended ssDNA that is bound by Rad51 or Dmc1, then force exerted on the phosphodiester backbone between base triplets would be on the order of 20-30 piconewtons [54]. Making the same assumptions, the force needed to extend the phosphodiester backbone to match the extended state of the ADP-bound forms of the Rad51 or Dmc1 filaments would be on the order of ∼100 pN [54]. Although the extension of the ssDNA within a Rad51 or Dmc1 filament is certainly not isotropic, these simple comparisons serve to illustrate that the increased extension on the ssDNA that would be needed to match to extension of the ADP-bound state would necessarily involve exerting increasing forces on the ssDNA which may in turn contribute to destabilizing the binding interactions.

**Figure 6.**
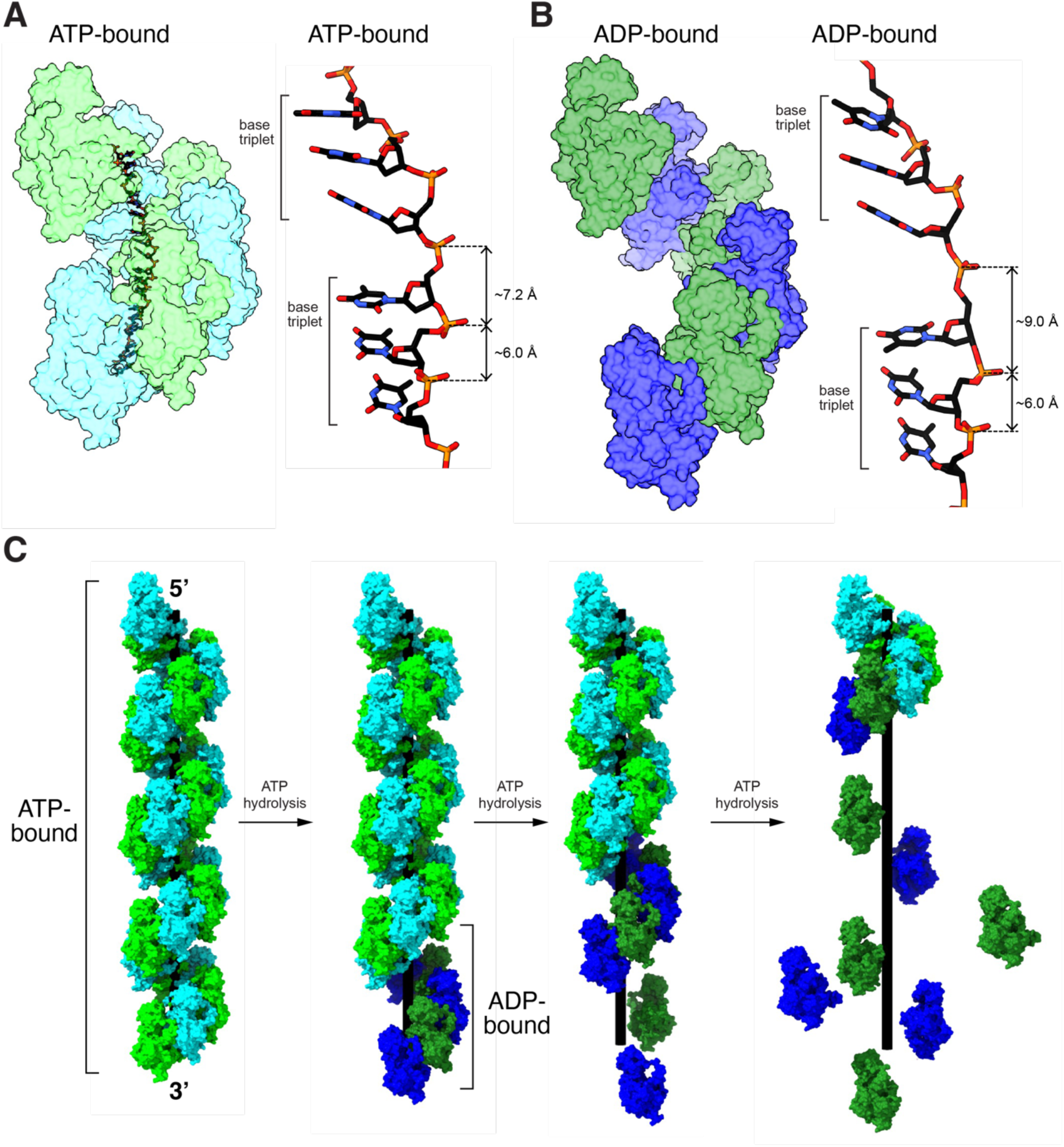
Model for Rad51 and Dmc1 nucleoprotein filament disassembly. **(A)** Rad51 and Dmc1 in the ATP-bound states give exhibit extended nucleoprotein filaments and the bound ssDNA is organized into base triplets with an axial rise of ∼7.2 Å between the last base of one triplet and the first base of the next triplet. **(B)** Conversion to the ADP-bound state results in nucleoprotein filament length, which would require even greater extension of the phosphate backbone in order the base triples to maintain correct register with the protein protomers. **(C)** The nucleoprotein filaments begin in the ATP-bound state and conversion of end-bound protomers leads to loss of contacts with the ssDNA and disruption of the protein-protein interfaces leading to protein dissociation from the ssDNA. Successive rounds of ATP hydrolysis can lead to complete disassembly of the nucleoprotein filaments.

Given these considerations, one potential model for how the ATP hydrolysis cycle is coupled to nucleoprotein filament disassembly is that conversion of the end-bound protein monomers, or small clusters of end-bound proteins (*e.g.* 1-3 monomers) [39] to the ADP-bound state is coupled to entry of these monomers into a highly extended configuration that is no longer compatible with ssDNA binding due to the increased tension that would be required to maintain the DNA in such a highly extended state (Figure 6C). Conversion of the ADP-bound state also coincides with structural changes in the protein-protein interfaces between proteins within the filament which likely favor dissociation of these protomers from the ends of the nucleoprotein filaments (Figure 6C) and loss of the second metal ion, as previously described [45].

In considering our model of ATP-hydrolysis coupled structural changes and nucleoprotein filament disassembly it is important to highlight one caveat, namely, we are not assembling filaments with ATP or allowing them to go through the normal process of ATP hydrolysis. Instead, we are preparing the filaments in the ADP-bound state by providing ADP as the only nucleotide cofactor within the sample mixture. We cannot rule out the possibility that filaments undergoing the normal hydrolysis of ATP to ADP plus Pi may be structurally distinct from those that are assembled in the presence of ADP alone.

## AKNOWLEDGMENTS

We thank members of the Greene laboratory for carefully reading this manuscript. This research was support by NIH Grant R35GM118026 (to E.C.G.). S.K. was partially supported by the Columbia University Summer Undergraduate Research Fellowships (SURF) program, the Columbia College Summer Funding Program, and an NSF Grant MCB-1817315 (to E.C.G.). The CryoEM was in part supported by the National Cancer Institute’s National Cryo-EM Facility at the Frederick National Laboratory for Cancer Research under contract 75N91019D00024, some of the work was performed at the National Center for CryoEM Access and Training (NCCAT), and the Simons Electron Microscopy Center located at the New York Structural Biology Center (NYSBC) and the Columbia University Cryo-Electron Microscopy Center operated by the NYSBC, both of which are supported by the NIH Common Fund Transformative High Resolution Cryo-Electron Microscopy program (U24 GM129539) and by grants from the Simons Foundation (SF349247) and NY State Assembly.

## AUTHOR CONTRIBUTIONS

Y.S. purified all proteins and conducted all CryoEM data collection and analysis and prepared all figures for publication. S.K. assisted with protein cloning and purification. E.C.G. assisted with data analysis and wrote the manuscript together with Y.S.

## METHODS

### Proteins purification

*S. cerevisiae* Rad51 was overexpressed in *E. coli* BL21 (DE3) Rosetta2 cells transformed with a plasmid encoding 6XHis-SUMO-Rad51. Cells were grown in 2L LB media containing 100 μg/ml carbenicillin and 35 μg/ml chloramphenicol at 37°C to an OD_600_ = 0.6, then induced with 0.5 mM IPTG and grown for 3 hours at 37°C. Cells were harvested by centrifugation and the resulting cell paste was resuspended in 50 ml lysis buffer (50 mM Tris-HCl [pH 7.5], 10% glycerol, 1M NaCl, 1mM DTT, 0.5 mM PMSF, 15 mM imidazole, 0.1% Tween 80, 1 protease inhibitor tablet [Roche, Cat No: 5892970001]) and then lysed by sonication. The lysate was centrifuged at 35,000 rpm for 45 mins followed by precipitating the supernatant with 12g ammonium sulfate for 1 hour. The precipitate was spun down at 10,000 rpm for 30 mins. The ammonium sulfate pellet was dissolved in 50 ml binding buffer (25 mM Tris-HCl [pH 7.5], 10% glycerol, 200 mM NaCl, 0.1% Triton X-100, 15 mM imidazole, 5 mM beta-mercaptoethanol) and applied to a 5 ml HisPur^TM^ Ni-NTA resin (Thermo Fisher Scientific) equilibrated with the same binding buffer. The protein was eluted with 10 ml elution buffer (25 mM Tris-HCl [pH 7.5], 10% glycerol, 200 mM NaCl, 200 mM imidazole, 0.1% Triton X-100). SUMO protease was added to the elution, followed by dialyzing for 16 hours at 4°C in dialysis buffer (50 mM Tris-HCl [pH 7.5], 200 mM NaCl, 10% glycerol, 15 mM imidazole, 1 mM DTT). The sample was re-applied to the 5 ml Ni-NTA resin equilibrated with the binding buffer and the flow-through was collected and concentrated to 80 μM using a spin concentrator (Vivaspin 6, 10 kDa MWCO; Cytiva, Cat No: 28932296). The protein was flash-frozen in liquid nitrogen and stored at –80°C.

*S. cerevisiae* Dmc1 was overexpressed in *E. coli* BL21 (DE3) Rosetta2 cells transformed with a plasmid encoding 6XHisDmc1. Cells were grown in 2L LB media containing 100 μg/ml carbenicillin and 35 μg/ml chloramphenicol at 37°C to an OD_600_ = 0.8, then induced with 0.1 mM IPTG and grown for 16 hours at 16°C. The cells were harvested by centrifugation and the cell paste was suspended in 100 ml lysis buffer (50 mM Tris-HCl [pH 7.5], 10% glycerol, 500 mM KCl, 0.01% Triton X-100, 1 mM DTT, 2 mM ATP, 2 mM MgCl_2_, 1 mM PMSF, 1 protease inhibitor tablet [Roche, Cat No: 5892970001]) and then lysed by sonication. The lysate was centrifuged at 35,000 rpm for 45 mins and the supernatant was applied to 5 ml Talon resin (Takara) equilibrated with the binding buffer (25 mM Tris-HCl [pH 7.5], 10% glycerol, 150 mM KCl, 0.01% Triton X-100, 2 mM ATP, 2 mM MgCl_2_). The resin was washed with wash buffer (25 mM Tris-HCl [pH7.5], 10% glycerol, 500 mM KCl, 0.01% Triton X-100, 2 mM ATP, 2 mM MgCl_2_) followed by re-equilibrating with the binding buffer. The protein was eluted with 10 ml elution buffer (25 mM Tris-HCl [pH 7.5], 10% glycerol, 150 mM KCl, 200 mM imidazole, 0.01% Triton X-100, 2 mM ATP, 2 mM MgCl_2_) followed by dialyzing for 16 hours at 4°C in dialysis buffer (25 mM Tris-HCl [pH 7.5], 100 mM KCl, 10% glycerol, 0.01% Triton X-100, 2 mM MgCl_2_, 0.5 mM EDTA). After dialysis, the sample was injected to a 1 ml heparin sepharose column (GE Healthcare) and fractionated with a 100 – 600 mM KCl gradient. The fractions containing Dmc1 were combined, concentrated to 80 μM with a spin concentrator (Vivaspin 6, 10 kDa MWCO; Cytiva, Cat No: 28932296) then flash-frozen in liquid nitrogen and stored at –80°C.

### CryoEM sample preparation

To prepare the *S. cerevisiae* Rad51 filaments in the ADP-bound state for CryoEM analysis, purified Rad51 (20 μM) was mixed with 0.5 µM of a 96–mer ssDNA (IDT; 5’– AAT TCT CAT TTT ACT TAC CGG ACG CTA TTA GCA GTG AAA ATT TCC TGA TAG TCG TCA CCG CGT TTT GCG CAC TCT TTC TCG TAG GTA CTC AGT CCG–3’) in HR buffer (30 mM HEPES [pH 7.5], 50 mM KCl, 20 mM MgCl_2_, 1 mM DTT) supplemented with 5 mM ADP and incubated at 30°C for 10 minutes. A sample volume of 3.5 µl was applied to a glow-discharged UltrAuFoil grid (R 0.6/1, 300 mesh), blotted for 4.5 seconds and plunge-frozen in liquid ethane using Vitrobot Mark IV (FEI, USA) at 100% humidity and 4°C.

To prepare the *S. cerevisiae* Dmc1 filaments in the ADP-bound state for CryoEM analysis, purified Dmc1 (2.5 μM) was mixed with 0.125 µM of 96–mer ssDNA in HR buffer (as above) supplemented with 1.5 mM CaCl_2_ and 5 mM ADP. The sample was incubated at 30°C for 5 minutes. The samples were then supplemented with 8 mM CHAPSO (3-([3-cholamidopropyl]dimethyammonio)-2-hydroxy-1-propanesulfonate; Hampton Research, Cat No: HR2-406-87), prior to being applied to UltrAuFoil grid (R 0.6/1, 300 mesh) and plunge-frozen in liquid ethane, as described above for Rad51.

### CryoEM data acquisition

Samples were initially screened using a Glacios (Thermo Fisher, 200 keV) at Columbia University Irving Medical Center. Grids selected for high-resolution data collection were imaged using a Titan Krios (Thermo Fisher) microscope equipped with a K3 direct electron detector (Gatan). Rad51 data was collected at National Cryo-Electron Microscopy Facility at National Cancer Institute with the microscope operated at 300 keV in electron counting mode, with a defocus range of −0.75 μm to −1.75 μm and magnification of 105,000x corresponding to 0.855 Å image pixel and a nominal dose of 50 e^-^/Å^2^. A total of 14,085 micrographs were collected. Dmc1 data was collected at Columbia University Irving Medical Center with the microscope operated at 300 keV in electron counting mode, with a defocus range of −0.75 μm to −1.75 μm and magnification of 105,000x corresponding to 0.823 Å image pixel size and a nominal dose of 58 e^-^/Å^2^. Total of 6,390 micrographs were collected.

### Rad51-ADP cryoEM data processing

Raw movies were processed using cryoSPARC v4.3.1 [55]. The beam induced motion was corrected and CTF was estimated by patch motion correction and patch CTF estimation jobs [56, 57]. The aligned micrographs were manually examined for quality such as ice contamination, and 12,982 micrographs were selected for further processing (Figure S1A). 1,748,800 particles were picked by template-free blob picking and extracted with a box size of 360×360. After removing junk particles, such as denatured proteins and low-quality filaments, by three rounds of 2D classification, 615,261 particles were used to generate 4 ab initio 3D templates followed by heterogenous refinement (Figure S1B-C). A class corresponding to the nucleoprotein filament with 428,741 particles was subjected for another round of 2D classification, clean-up resulted in 330,601 final particles representing the Rad51-ADP density map (Figure S1D). The density map was further polished by non-uniform refinement and CTF refinement. The nominal resolution of the density map of 3.37 Å was estimated by 0.143 gold standard Fourier Shell Correlation (FSC) cut off (Figure S1E).

### Dmc1-ADP cryoEM data processing

Raw movies were processed using cryoSPARC v4.3.1 [55]. The beam induced motion was corrected and CTF was estimated by patch motion correction and patch CTF estimation jobs [56, 57]. The aligned micrographs were manually examined for quality such as ice contamination, and 6,044 micrographs were selected for downstream processing (Figure S2A). 1,184,484 particles were picked by template-free blob picking and extracted with a box size 360×360. During the imaging processing, we noticed that the NPF showed bundled filaments in the micrographs and 2D class average images (Figure S2B). After 2D classification, 1,124,382 particles were selected for generating 4 ab initio 3D templates followed by heterogenous refinement (Figure S2C). A class with 960,233 particles corresponding to nucleoprotein filament was examined for possible contacts between the bundled filaments, but no clear atomic contacts between the filaments were identified. Moreover, the same exact symmetry was observed between the filaments suggesting each was identical. Therefore, we applied a focus mask corresponding to a filament and then used this mask to generate a single filament map (Figure S2D). Further particle subtraction and local refinement of the filament, resulting in the nominal resolution at 2.74 Å estimated by 0.143 gold standard FSC cut off (Figure S2E).

### Structure Refinement

To refine the Rad51-ADP and Dmc1-ADP filament structures, our previously determined cryoEM structures of Rad51-ATP bound state (PDB: 9D46) and Dmc1-ATP bound state (PDB: 9D4N) were used as starting models [44]. Each protomer from the ATP-bound state structures were treated as a separate monomer and fitted into the corresponding cryoEM density maps using ChimeraX [58]. After initial rigid-body refinement in Phenix [59], amino acid residue sidechains were manually inspected and corrected for fitting into the density map using Coot [60]. The ssDNA and disordered part of the DNA binding loops were not modeled in the structures as the density is disordered. Fitted models were real space refined with the secondary structure and Ramachandran restraints in Phenix [59].

### Structure Analysis

The pitch and monomers per turn values were analyzed by symmetry search utility function of cryoSPARC with search parameters of helical pitch between 10 Å and 150 Å and number of subunits per turn between 5 and 8 [55]. The best local minima values are reported. Overall protomer-protomer contacts were initially identified by using Mapiya web service [61]. The interactions were further investigated by contacts function in ChimeraX [58]. All the interactions were verified by visual inspection. All the figures were generated by ChimeraX.

## Data Availability

The refined structural models and cryo-EM density maps have been deposited in the Protein Data Bank (www.rcsb.org) and Electron Microscopy Databank (www.ebi.ac.uk/emdb) under accession numbers PDB:9NJK and EMD-49485 for Rad51- ADP and PDB:9NJR and EMD-49488 for Dmc1-ADP.

---

**Table S1.**
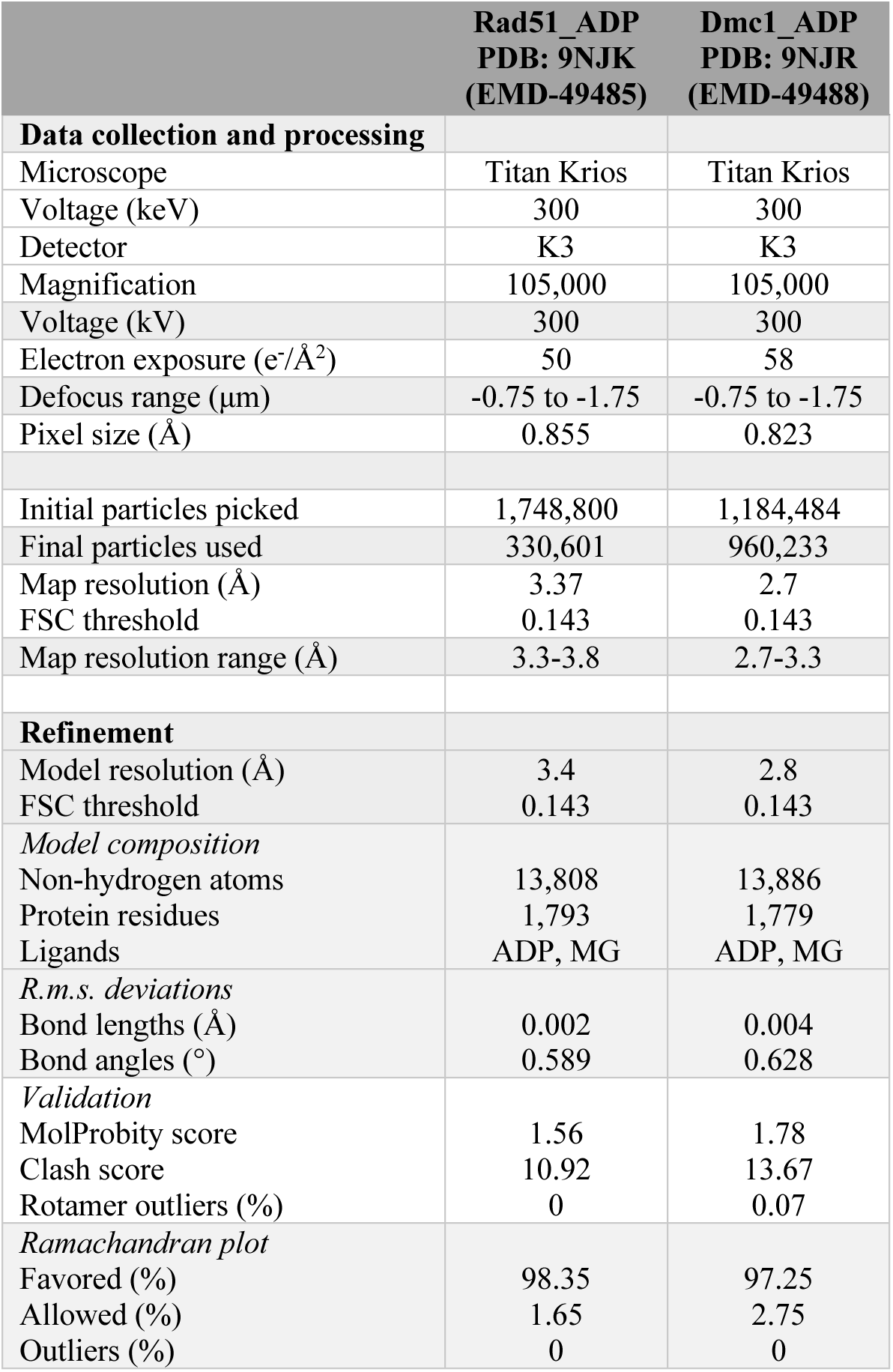
CryoEM parameters.

**Figure S1.**
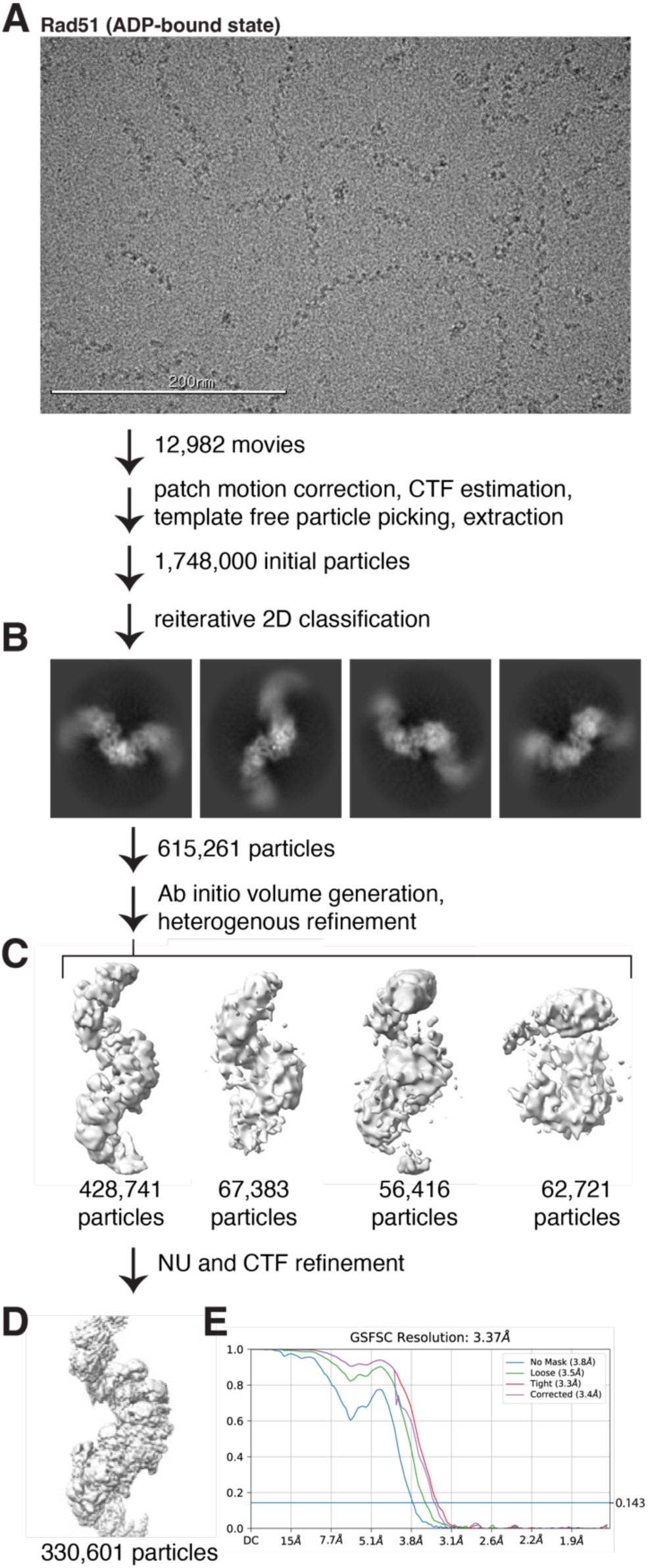
Cryo-EM image processing pipeline of the Rad51-ADP nucleoprotein filament. **(A)** Representative micrograph used for the Rad51-ADP nucleoprotein filament data processing. **(B)** Representative 2D classes selected for 3D classification. **(C)** 3D map generation and refinements. **(D)** Final 3D map reconstruction for the Rad51-ADP nucleoprotein filament. **(E)** Fourier shell correlation curve of the final electron density map.

**Figure S2.**
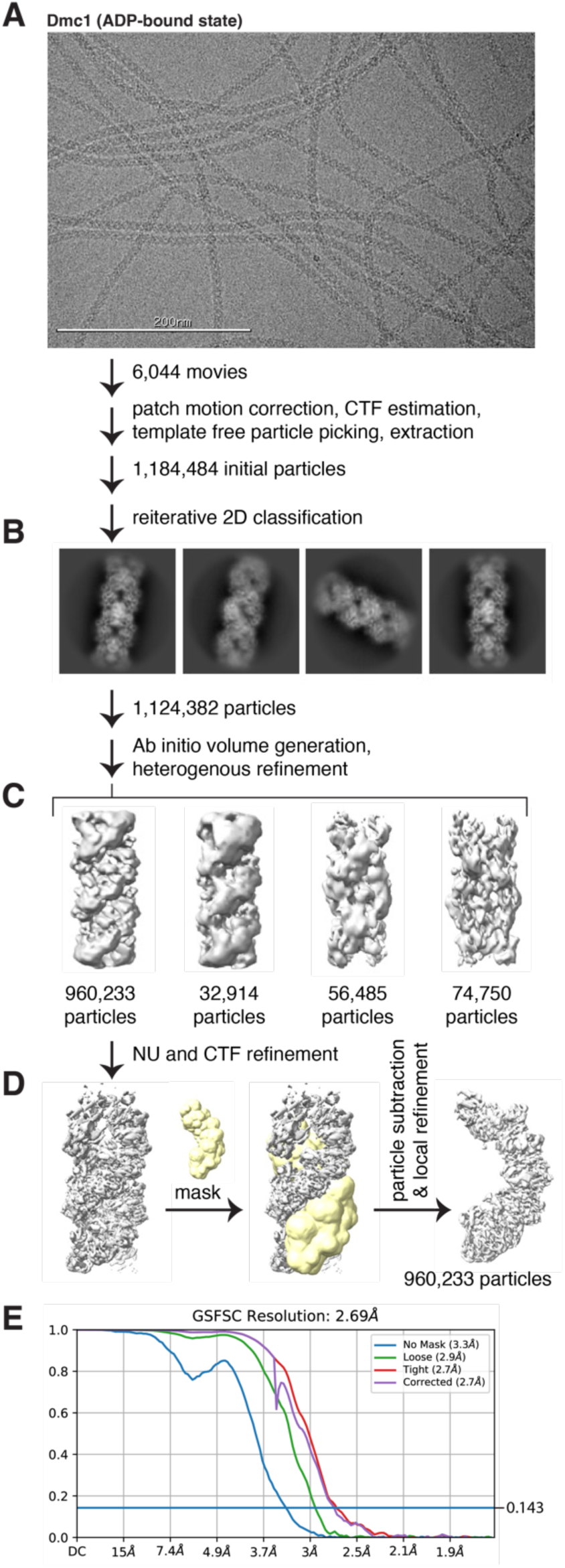
Cryo-EM image processing pipeline of the Dmc1-ADP nucleoprotein filament. **(A)** Representative micrograph used for the Dmc1-ADP nucleoprotein filament data processing. **(B)** Representative 2D classes selected for 3D classification. **(C)** 3D map generation and refinements. **(D)** Final 3D map reconstruction for the Dmc1-ADP nucleoprotein filament. **(E)** Fourier shell correlation curve of the final electron density map.

**Figure S3.**
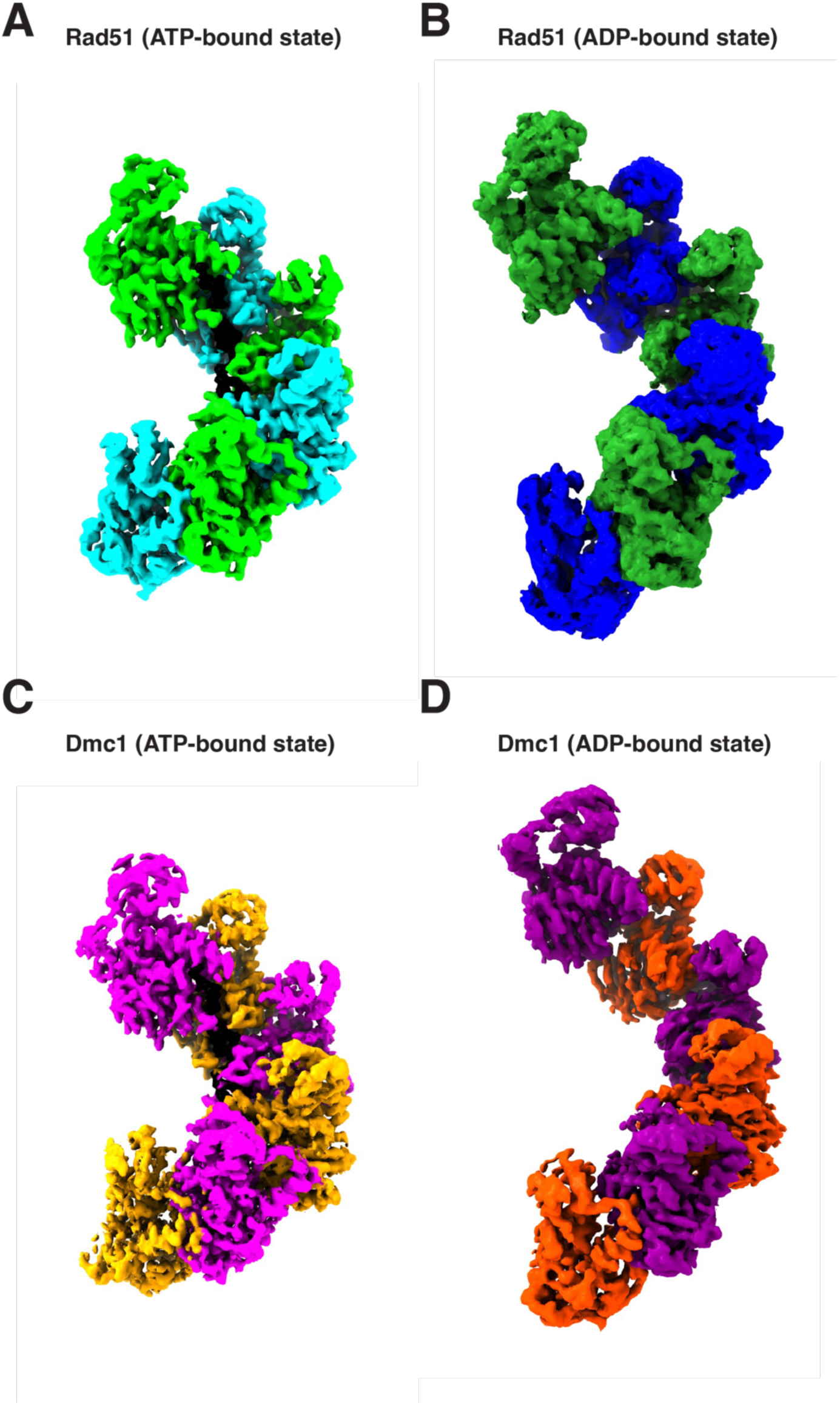
Cryo-EM density maps for the ATP- and ADP-bound nucleoprotein filaments. (A) Rad51 in the ATP bound state. (B) Rad51 in the ADP-bound state. (C) Dmc1 in the ATP bound state. (D) Dmc1 in the ADP-bound state. For each panel, a subsection of the nucleoprotein comprised of six Rad51 or Dmc1 monomers is shown, and the different protein monomers are highlighted in alternating colors. The ssDNA is shown in black.

**Figure S4.**
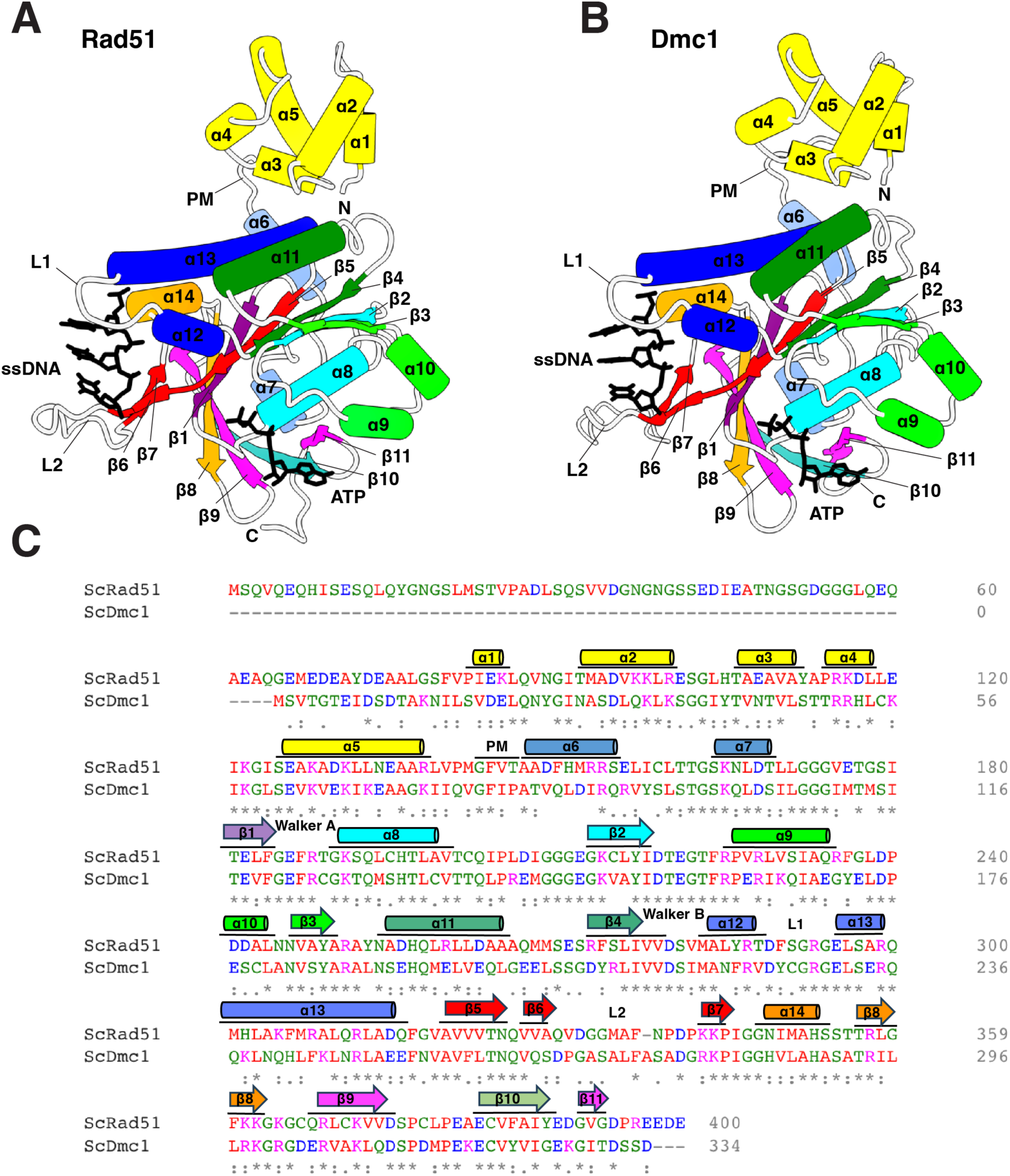
Diagrams of Rad51 and Dmc1 secondary structure topology. **(A)** Ribbon diagram of *S. cerevisiae* Rad51 in the ATP-bound state (PDB ID: 9D46)[44] showing the number designation for each alpha helix and beta strand. Also highlighted are the Walker A and B motifs, DNA binding loops L1 and L2, and the FxxA polymerization motif (PM). The bound ssDNA and ATP are shown in black. **(B)** Ribbon diagram of *S. cerevisiae* Dmc1 in the ATP-bound state (PDB ID: 9D4N)[44] showing the number designation for each alpha helix and beta strand. **(C)** Sequence alignment of *S. cerevisiae* Rad51 (UniProt ID: P25454) and Dmc1(UniProt ID: P25453) (residues in red are small, hydrophobic, or both [A, V, I, L, M, F, P]; residues in blue have acidic side chains [D, E]; residues in magenta have basic side chains [K, R]; green residues correspond to all others [H, S, T, N, Q, C, G, Y, W]; “**\***”indicates identical residues in Rad51 and Dmc1; “.” indicates residues have weakly similar properties; “:” indicates strongly similar residues). The sequence was aligned using Clustal Omega [62].

**Figure S5.**
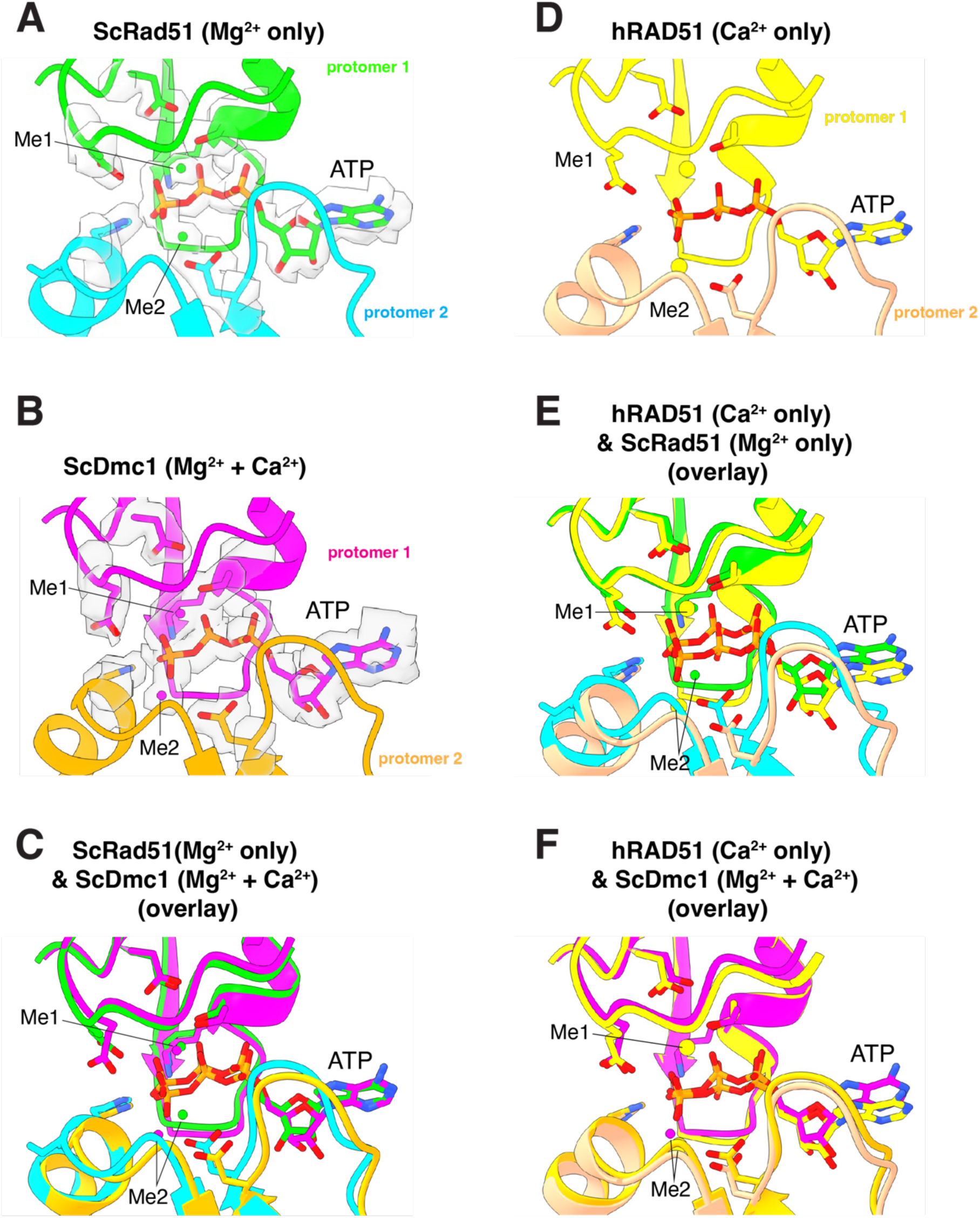
Differences in second metal ion positioning for samples containing Ca^2+^. **(A)** ATP-binding pocket of *S. cerevisiae* Rad51 highlighting the positions of the two metal ions (PDB ID: 9D46)[44]; the sample was prepared with 20 mM Mg^2+^. **(B)** ATP-binding pocket of *S. cerevisiae* Dmc1 highlighting the positions of the two metal ions (PDB ID: 9D4N)[44]; the sample was prepared with 20 mM Mg^2+^ plus 1.5 mM Ca^2+^. **(C)** Overlay of *S. cerevisiae* Rad51 and Dmc1 highlighting the difference in position of the second divalent metal ion. **(D)** ATP-binding pocket of human RAD51 highlighting the positions of the two metal ions (PDB ID: 8BQ2)[45]; the sample was prepared with 5 mM Ca^2+^. **(E)** Overlay of the ATP-binding pocket from *S. cerevisiae* Rad51 [44] and human RAD51 (PDB ID: 8BQ2)[45]. **(F)** Overlay of the ATP-binding pocket from *S. cerevisiae* Dmc1 [44] and human RAD51 (PDB ID: 8BQ2)[45]. In **(A-F)** Me1 and Me2 are used denote the first and second metal ion binding sites.

**Figure S6.**
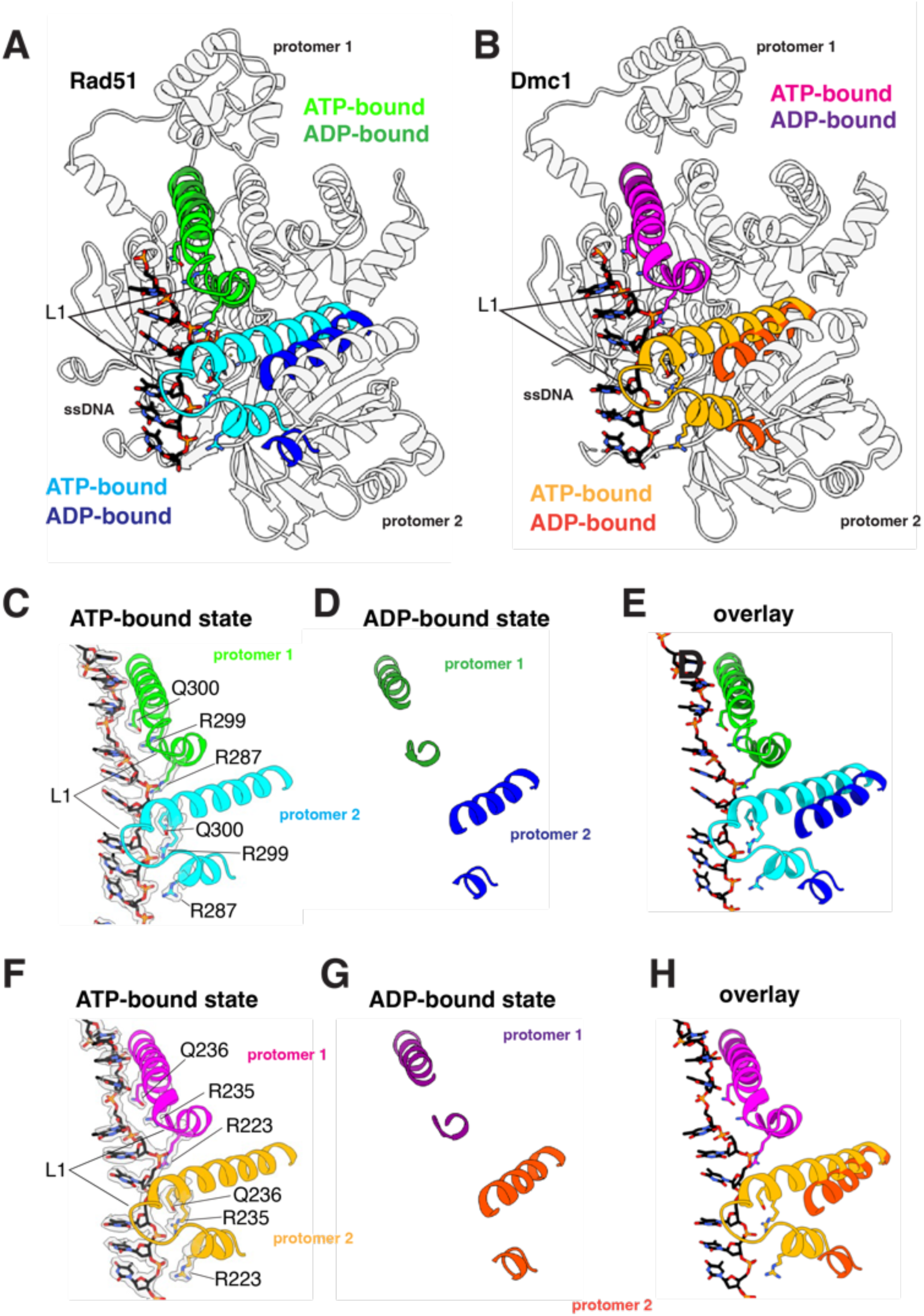
Loss of L1 contacts with ssDNA in the ADP-bound state. **(A)** Overlay of two adjacent Rad51 protomers in the ATP- and ADP-bound states, as indicated, highlighting the location of the L1 DNA-binding loop, α helix 12 and α helix 13. **(B)** Overlay of two adjacent Dmc1 protomers in the ATP- and ADP-bound states, as indicated, highlighting the location of the L1 DNA-binding loop, α helix 12 and α helix 13. **(C)** Close-up view of the Rad51 L1 contacts with the bound ssDNA substrate in the ATP-bound state. **(D)** Close-up view showing the loss of Rad51 L1 contacts with the bound ssDNA substrate in the ADP-bound state. **(E)** Overlay of the Rad51 region encompassing the L1 DNA-binding loop, α helix 12 and α helix 13 in the ATP- and ADP-bound states. **(F)** Close-up view of the Dmc1 L1 contacts with the bound ssDNA substrate in the ATP-bound state. **(G)** Close-up view showing the loss of Dmc1 L1 contacts with the bound ssDNA substrate in the ADP-bound state. **(H)** Overlay of the Dmc1 region encompassing the L1 DNA-binding loop, α helix 12 and α helix 13 in the ATP- and ADP-bound states.

